# Genetic and environmental factors regulating soybean reproductive stages and their transitions

**DOI:** 10.1101/2025.09.10.675456

**Authors:** Swapan Chakrabarty, Johnathon M. Shook, Asheesh K. Singh

**Affiliations:** Department of Agronomy, Iowa State University, Ames, IA 50010

**Keywords:** complex trait, reproductive maturity, genetic effect, photoperiod, thermal accumulation, maturity model

## Abstract

The reproductive stage of soybean is influenced by the effect of genotype, environment, and their interactions. While days to flowering and days to full maturity have been studied, a systematic and comprehensive study that investigates the variation in days to each stage and the role of maturity-related genes and environmental variables is lacking. Therefore, we studied 508 unique accessions from the USDA germplasm collection from maturity group 0-IV, and a set of 67 near-isogenic lines differing for maturity-related genes. Field experiments and evaluations were conducted in central Iowa, USA. The days to each of the reproductive stages, R1-R8, were recorded. We report considerable variation in the duration of reproductive growth stages between flowering and maturity, which is largely explainable by known flowering and maturity genes as well as environmental variables, day length, and growing degree days. Besides the known maturity-related genes *E1*, *E2*, and *Dt1*, we identified two novel SNPs, such as *Glyma.01G180600* and *Glyma.10G221300*, as potential targets for genetic regulation of reproductive stages. We also captured two other loci, *Glyma.08G216800* and *Glyma.04G088100* for day length and growing degree days, respectively, that revealed dynamic regulation of environmental gradients on the reproductive stages. Furthermore, we developed a random forest-based genetic maturity model that can predict genetic and environmental effects across a wide range of genotypes. This study broadens the understanding of the factors that contribute to reproductive development, which will help to develop cultivars that combine the optimal combinations of stage durations for a higher seed yield and enhanced resilience.

## 1 INTRODUCTION

As with many crop species, soybean phenology is commonly characterized based on easily measurable morphological characteristics using the staging code developed by Fehr and Caviness 1977 (1). During the vegetative stages, soybean is characterized by the number of nodes with unfurled leaves, which develop from the first node above the cotyledons and up. In the vegetative period, soybeans grow very rapidly and devote their energy to leaf, stem, and root tissue development. The initiation of flowering signals indicates the transition to the reproductive stage. Eight reproductive stages (R1 to R8) are generally recognized: two stages of flower initiation, two stages of pod formation and elongation, two stages of seed formation and expansion, and two stages of physiological maturity (Figure 1A). The growth and development aspects are directly related to the farmer’s decisions for the choice of cultivar, time of planting, and days to maturity for harvesting the crop. The decision-making process is further complicated by the weather (2). Stresses and injuries during the reproductive phases can cause significant seed yield loss, as time and resources for compensatory growth are limited (3,4).

**Figure 1.**
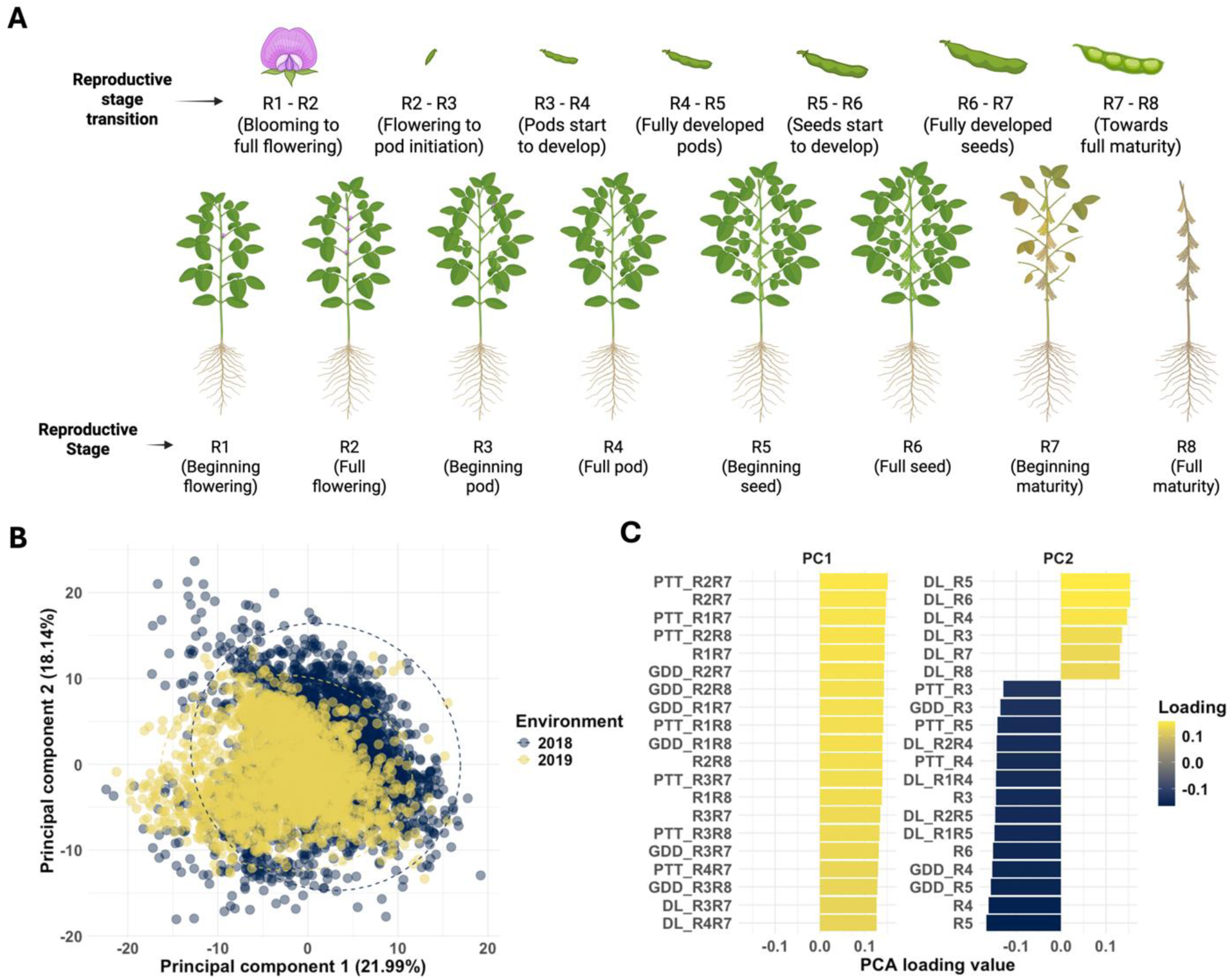
Phenotypic variations during reproductive stages and their transitions in a soybean diversity panel. (A) Depicting different reproductive stages (R1-R8) and their transitions (R1R8-R7R8), (B) PCA scatter plot with 95% confidence ellipses indicating genotype distributions for 2018 (dark blue) and 2019 (yellow), and (C) Top contributors to principal component 1 (PC1) and principal component 1 (PC2) are shown in bar plot form. The variables PTT, GDD, and DL indicate photothermal time, growth degree days, and day length, respectively. The contributions of each trait to the primary axis are reflected in the loadings. Part of this figure was created in https://BioRender.com.

One of the most important traits in soybean breeding and production is seed yield (5). Manipulation of the soybean developmental timeline holds potential for yield improvement through different mechanisms. Ideotype breeding attempts to optimize the combination of traits to achieve higher yield through physiological and morphological changes (6). The timing of reproductive stages may be altered to achieve a more optimal source-sink relationship and to minimize source limitation. For example, a prolonged flowering period may result in greater pod set, as fewer flowers would be competing for resources at any time (7). The developmental stages and their duration can be altered to avoid abiotic and biotic stresses (8). For example, if biotic or abiotic stresses are likely at a specific time of year, such as due to weather patterns or insect reproduction cycles, one way to mitigate or lessen the negative impact is to alter the timing of developmental stages to minimize the risk of these stresses coinciding with the most susceptible stages. Another farming strategy is to manipulate the planting date, which has cascading effects on soybean phenology. Among the reproductive stages, R1 (Flowering), R4 (Pod initiation), and R8 (physiological maturity) are more commonly studied (1,9); however, there is a lack of literature on comprehensive reproductive stage studies. To optimize soybean maturity and production, it is essential to comprehend how genetics and the environmental variables regulate reproductive development. Understanding the mechanism underlying phenotypic variation requires determining the extent to which populations exhibit underlying genetic variation, and the effect of the environmental variables in natural field conditions (10,11). However, it is challenging to predict responses to variable climates without knowing the particular environmental cues that cause plastic responses, including in more extreme and less studied climates (12). Early planting in the U.S. Midwest region is a recommended management strategy to boost soybean yield (13), but personnel, equipment, logistics, and environmental issues might cause planting to be delayed. Soybeans are more vulnerable to spring killing frost, early-season insects, seedling disease, and damaging rainfall events that could lead to a substandard emergence when early planting is feasible (14). In the MGII and earlier maturities, the risk of frost rises and the time to emergence falls with delayed planting. Diseases of the seedlings and roots are less likely to occur when planting and emergence times are shorter (2). Therefore, studies on soybean maturities, the effect of known maturity-related genes, the usefulness of identifying new genetic factors, the relationship of genetics with the reproductive stages, and their transitions are crucial.

The understanding of the genetic underpinnings of developmental stage timing is further complicated by both temperature and photoperiod, as they are known to affect the timing of reproductive stages (15–18). The findings of a simulation study by Forecast and Assessment of Cropping sysTemS (FACTS) indicated that temperature affects early vegetative development (days from planting to blooming) more than photoperiod, and this effect is especially noticeable in the maturity group (MGII). Temperature was significant in the maturity group MGIV, but the effect of photoperiod (influenced by latitude) was greater in the MGII (2). The genetic control of flowering (R1) and maturity (R8) timing has been extensively studied in soybean due to its effect on adaptation to a specific environment (19–22). At least 14 genetic loci have been identified that have a clear effect on the latitudinal adaptability of a given soybean variety in the field (Supplementary File 1 - Table S1). These include eleven *E* genes (*E1*-*E11*) and three additional genes (*Dt1*, *Dt2*, *J*), which cause major shifts in latitudinal adaptability (between 5-30 days to maturity per *E* locus) (23,24). *E1*, a B3 transcription factor, epistatically controls time to maturity through its interactions with several other *E* genes (25). In the presence of epistasis, the allelic effect of *E1* to *e1* delays maturity by nearly three weeks. *E2* is an ortholog of the Arabidopsis gene *GIGANTEA* (26). *E3* and *E4* are phytochrome A genes (27,28). *Dt1* is an ortholog of Arabidopsis *TERMINAL FLOWER1* (29); while *Dt2* is a MADS-domain factor gene (30). These are the commonly included genes in soybean maturity models, as these genes have been mapped in multiple studies, allele-specific markers have been developed, and genes have been cloned (26–31). Eight (*E1*-*E4*, *E9*, *Dt1*, *Dt2*, and *J*) of these 14 genes have the causal gene identified, with the corresponding gene sequence available from the Williams 82 reference genome of soybean (32).

While the role of these genes on days to flowering and days to maturity has been studied (22,33,34), minimal information is available on the effect of these genes during different reproductive stages. In a quest for further information on the genetic control of developmental stages, several genome-wide association studies have previously reported on flowering and/or maturity timing (35–37). As mentioned previously, there is a lack of information on intermediate stages between R1 and R8 (38). Additionally, as plant breeders have traditionally worked with a narrow genetic base (39,40), a smaller repertoire of diversity within development-related genes in the germplasm pool, insufficient information is available about the genetics of maturity-influencing genes, growth stage durations, and genetic control. In this study, we studied a large panel of diverse soybean genotypes and a panel of near-isogenic lines (NILs) that were grown in central Iowa for two years (2018 and 2019) and investigated the genetic variation for eight reproductive growth stages, and the effect of environmental variables on different reproductive stages (R1-R8). The objectives of this work were to: (1) investigate the genetic variation for reproductive growth stages in soybean, (2) perform genome-wide association analysis on reproductive growth stages, and study the effect of *E*, *J*, and *Dt* loci on reproductive growth stages, and (3) model the effect of genetic loci and environmental variables influencing reproductive growth stages.

## 2 MATERIALS AND METHODS

### 2.1 Plant material and experimental design

A diversity panel including SoyNAM parents and landrace accessions from the core collection was grown in 2018 (42°00’35.5“N, 93°44’01.9“W) and 2019 (42°01’9.32”N, 93°46’11.5”W) near Ames, IA, and was used in this study. In 2018, 508 lines were included within this study. The accessions that were viny-type, agronomically inviable, or did not produce sufficient seed in the winter nursery increase were removed from the experiment grown in 2019. Subsequently, 450 genotypes were included in the 2019 field study. The full list of genotypes grown in each year is provided in Supplementary File 2 - Table S1. In 2018 and 2019, a concurrent experiment that included 67 near-isogenic lines (NILs) for 14 maturity-related genes was planted in the same field (Supplementary File 2 - Table S2). These NILs were primarily designed in Clark and Harosoy backgrounds, with allele swapping for one or more of the known maturity or stem termination genes (23).

In 2018, the diversity panel (508 accessions) was planted on May 29th in an RCB design with 6 reps, each plot consisting of a hill plot with three seeds planted per hill in a 30” x 30” grid. A single, representative plant was harvested for each genotype at full maturity, and hand-shelled for winter increase in Chile to minimize heterogeneity within accessions. Two pods from each of these plants were stored for genotyping. In 2019, 450 genotypes were planted on June 4th in an RCBD with 5 reps, each plot consisting of 7’ long paired rows on 30” rows. For the 67 near-isogenic line (NIL) panel, the plots were organized in an RCB design blocked by planting date, with three planting dates per year, staggered approximately 10-14 days apart to allow for dissection of photoperiod and thermal effects on time to each reproductive stage. All plots (in both years) consisted of hill plots with three seeds per plot on a 30” x 30” grid. Planting dates in 2018 were: May 29, June 7, and June 21, while planting dates in 2019 were: June 4, June 14, and June 24. In the later analysis in this study, each year was considered as an environment.

### 2.2 Phenotyping and environmental variables

Throughout the growing season, the development stage data were recorded when each plot reached R1, R2, R3, R4, R5, R6, R7, and R8 stages. We calculated the duration of each reproductive stage transition, such as R1 to R8 (R1R8), R2 to R8 (R2R8), and so on (1). Each plot was phenotyped at least once every three days, and if needed, backdating was applied to plots that appeared to reach a growth stage one or two days prior to the rating date. There were four environmental variables used in this study: day length (DL), growing degree days (GDD), diurnal temperature range (DTR), and photothermal time (PTT). For estimating the environmental indexes, the Global Historical Climatology Network (GHCN) database at the National Oceanic and Atmospheric Administration (NOAA)’s National Centers for Environmental Information (NCEI) (https://www.ncdc.noaa.gov/) was used for retrieving daily maximum (T_max_) and minimum (T_min_) air temperature (Fahrenheit). DL was calculated by using the function daylength in the R package geosphere. GDDs were obtained by using the formula ((T_max_ + T_min_)/2 – T_base_), with T_max_ greater than 100°F adjusted to 100°F, T_min_ lower than 50°F adjusted to 50°F, and T_base_ considered as 50°F. The PTT was estimated as GDU x DL. The daily DTR was calculated by subtracting T_min_ from T_max_ (i.e., T_max_ – T_min_), without any adjustment.

### 2.3 Phenotypic variability and selection of environmental indices

For assessing the variation and effect of genotype, replication, and year (2018 and 2019) on the phenotype, we performed analysis of variance (ANOVA) for each reproductive stage, and their transitions, as well as their respective environmental variables such as DL, GDD, DTR, and PTT for individual years and combined. In the combined analysis, each year was considered as an environment. The coefficient of variation (CV) and Tukey’s honest significant difference (HSD) test were performed to evaluate the phenotypic differentiation among the genotypes. Further, we conducted principal component analysis (PCA) using the prcomp() function in R to examine multivariate patterns among variables and evaluate how environment and genetics contribute to phenotypic structure. For the identification of environmental variables that significantly contribute to the reproductive stages and their transitions, we used the means of DL, GDD, DTR, and PTT from the years (environment) 2018 and 2019. The environmental variables for each reproductive stage and their transitions were correlated with the respective reproductive stages and their transitions. The environmental variables that showed the most significant correlation with the reproductive stages and their transitions, as well as physiologically relevant, were used in further statistical analysis as environmental variables. All analyses were performed using R version 4.3.3 (41).

#### 2.3.1 Effect of genotype, environment, and their interaction

The best linear unbiased estimate (BLUE) of the genotype effect was estimated for each reproductive stage and their transitions using a linear model in the R package lme4 (42) and modelled as:

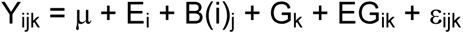

Where, μ indicates the mean of the population; E_i_, B(i)_j_, and G_k_ is the effect of environment (i.e., year), replication nested in environment, and genotype, respectively; EG_ik_ indicates the interaction between the environment and genotype; and e_ijk_ is the error.

The model considered the fixed effects of genotype, replication, environment, and genotype x environment (G x E). The model calculates each genotype’s mean trait performance across environments and the impacts of genotype, environment, and their interaction. After adjusting for replication and environmental effects, the genotype effect is the best linear unbiased estimate of each genotype’s performance across environments. To understand how genotypes respond to environmental variations, a joint regression analysis was performed (43,44). The environmental means were first estimated for each environment (2018 and 2019), and then the first environmental mean regression model (Finlay-Wilkinson Regression) (45) was fitted to each of the genotypes to assess the relation between the traits and the environmental mean, where the traits were modelled as a function of the environmental mean. The environmental mean regression model was described as:

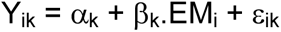

Where, Y_ik_ is the value of k in the environment i; α_k_ is the intercept (baseline performance of genotype k; β_k_ indicates the response of the genotype k to environment; EM_i_ indicates the mean trait value of environment i; and ε_ik_ indicates residual error In the second joint regression, a reaction norm was performed to understand the G x E interaction with the identified environmental variables. To investigate the genotype plasticity in response to the environmental variables, fitted values from the first regression were used, where the identified environmental variables replaced the environmental mean and were used as the explanatory variable. The model yielded the slope (plasticity) and intercept of the reaction norm.

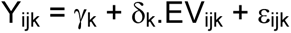

Where, Y_ijk_ is the fitted trait value for genotype k, environment i, replicate j from the BLUE model; γ_k_ indicates genotype-specific intercept; δ_k_ is the reaction norm plasticity slope in response to environmental variable (EV); EV_ijk_ is a scaled environmental variable; and ε_ijk_ is the residual error

Variance component analysis (VCA) was performed using a mixed-effect model applying the VCA() function in R to partition the variance of each reproductive stage and their transitions into components attributable to genotype, environment, replication, and their interaction (G x E). This analysis allows for a deeper understanding of the factors that contribute to the variability in the trait and guides breeding strategies. Furthermore, the magnitude of G x E interaction for each genotype was quantified by estimating the mean absolute residual from the BLUE model. This measure provided an estimate of the genotype’s performance across the environments. Based on the reaction norm and G x E magnitude of the genotypes to the specific environmental variables, they are classified into stable, plastic, and environment-dependent genotypes.

### 2.4 Genome-wide association studies (GWAS)

Genome-wide marker data for the diversity panel were downloaded from SoyBase using Wms82.a2.v2 as the reference genome (46). GWAS analysis was performed considering the attributable component for each of the reproductive stages and their transitions. In the GWAS analysis, we used BLUE estimates for capturing the SNP related to genotypic effect only; we also used reaction norm parameters, such as plasticity slope, to identify the SNPs related to the effect of the environmental variable, and intercept for identifying SNPs related to the average performance of the population across all the environments. GWAS was conducted using four models: FarmCPU (Random Model Circulating Probability Unification), BLINK (Bayesian-information and Linkage Disequilibrium Iteratively Nested Keyway), MLM (Mixed Linear Model), and MLMM (Multi-Locus Mixed Linear Model) (47–49), and run in R using the GAPIT v3 (Genome Association and Prediction Integrated Tool) R package (50). The GAPIT’s Bonferroni correction (0.05/number of SNPs) and a significance threshold of P-value 0.05 were used for multiple hypothesis testing. We used these four models to utilize their statistical and computational power for capturing SNPs related to known maturity-related genes and to identify novel candidate genes. In our study, the known maturity-related genes captured by any of these models were included; however, for the novel candidates, only the SNPs that were captured by more than one model were considered. The identified SNPs were annotated using the SoyBase database using Wms82.a2.v2 as the reference genome (32). We further performed a multi-step haplotype analysis utilizing the pegas package (51) in a custom R script to examine haplotype structure and linkage disequilibrium (LD) around important SNPs found through GWAS. It enabled the finding of local SNP blocks in strong LD with each GWAS SNP and described their haplotype patterns across the genotypes. To evaluate pairwise LD, we calculated the Pearson correlation coefficient with every other SNP in the dataset. SNPs that had an absolute correlation of at least 0.8 were considered in LD and added to a particular GWAS SNP’s haplotype block.

### 2.5 *De novo* marker analysis and development of a genetic maturity model

For *de novo* marker analysis, we conducted Kompetitive Allele Specific PCR (KASP) genotyping (LGC Genomics) based on previously reported variation within *E1*, *E2*, *E3*, *E4*, *E9*, *E10*, *Dt1*, *Dt2*, and *J*. Assays for twenty-two markers were developed and deployed across the near-isogenic lines (NILs) and diversity panel. Of these, ten markers showed no variation within either panel. The remaining markers (*E1*, *E2*, *E3*, *E4*, *Dt1*, *Dt2*, *J, E1E3*, *E1E4*, *E2E3*, *E2E4*, *and E3E4*), were included in the analyses. The *E1E3*, *E1E4*, *E2E3*, *E2E4*, *and E3E4* indicated the interactions between the loci that were not captured by individual loci. At first, we used the NIL panel for the prediction of the timing of the reproductive stages (R1-R8) through implementing a Random Forest regression framework using caret and randomForest packages in R. The dataset included genetic data such as genotype of known maturity and stem termination genes and interactions between E loci. The dataset also included days to reach each of the reproductive stages (R1-R8) and photothermal variables (DL and GDD) for each reproductive stage. In the model, the timing of the reproductive stages (R1-R8) was designated as the target variables. Using 10-fold cross-validation to ensure model robustness, we trained a distinct Random Forest regression model for each step (R1–R8). The caret package’s createDataPartition() was used to divide the data into training (80%) and testing (20%) subsets. The performance of each model was assessed using root mean square error (RMSE) and coefficient of determination (R^2^) of the test set after 500 trees (ntree = 500) were used for training. Further, the varImp() function was utilized to extract the feature importance from the training models.

## 3 RESULTS

### 3.1 Phenotypic variation of the genotypes for reproductive maturity

Phenotypic variation was higher for environment 2019 compared to 2018 within the diversity panel (Table 1, Supplementary File 1 - Figure S1). Earlier maturity groups reached each reproductive stage sooner than their late-maturing counterparts when examined as a whole (Table 1, Supplementary File 1 - Figure S1A & S1C). For transitioning to the reproductive stages, such as flowering to pod initiation (R1-R3), pod development to seed filling (R4-R6), and towards full maturity (R7-R8), the average time followed the same trend (Supplementary File 1 - Figure S1B & S1D, and Table S2). However, within maturity groups, there was still considerable variation. ANOVA results for individual and combined environments showed significant genotypic effects (p<0.05). In 2018 and 2019, the coefficient of variation (CV) was 9.15% and 8.90%, respectively (Supplementary File 2 - Table S3, S4 & S5). The CV in the combined dataset was 9.03%. We used PCA on standardized trait values from both environments to demonstrate multivariate variance. Of the overall variation, PC1 and PC2 accounted for 21.99% and 18.14%, respectively. A considerable divergence between the environment in 2018 and 2019 was shown by the PCA scatter plot (Figure 1B, and Supplementary File 2 - Table S6). Despite some overlap, the genotypes from 2018 and 2019 formed separate clusters that showed environmental differences. The top contributors to PC1 were the PTT and GDD, which were associated with later transitions, such as R2R7 and R1R8. On the other hand, PC2 was mostly driven by DL during earlier transitions, R3 to R5 (Figure 1C).

**Table 1.**
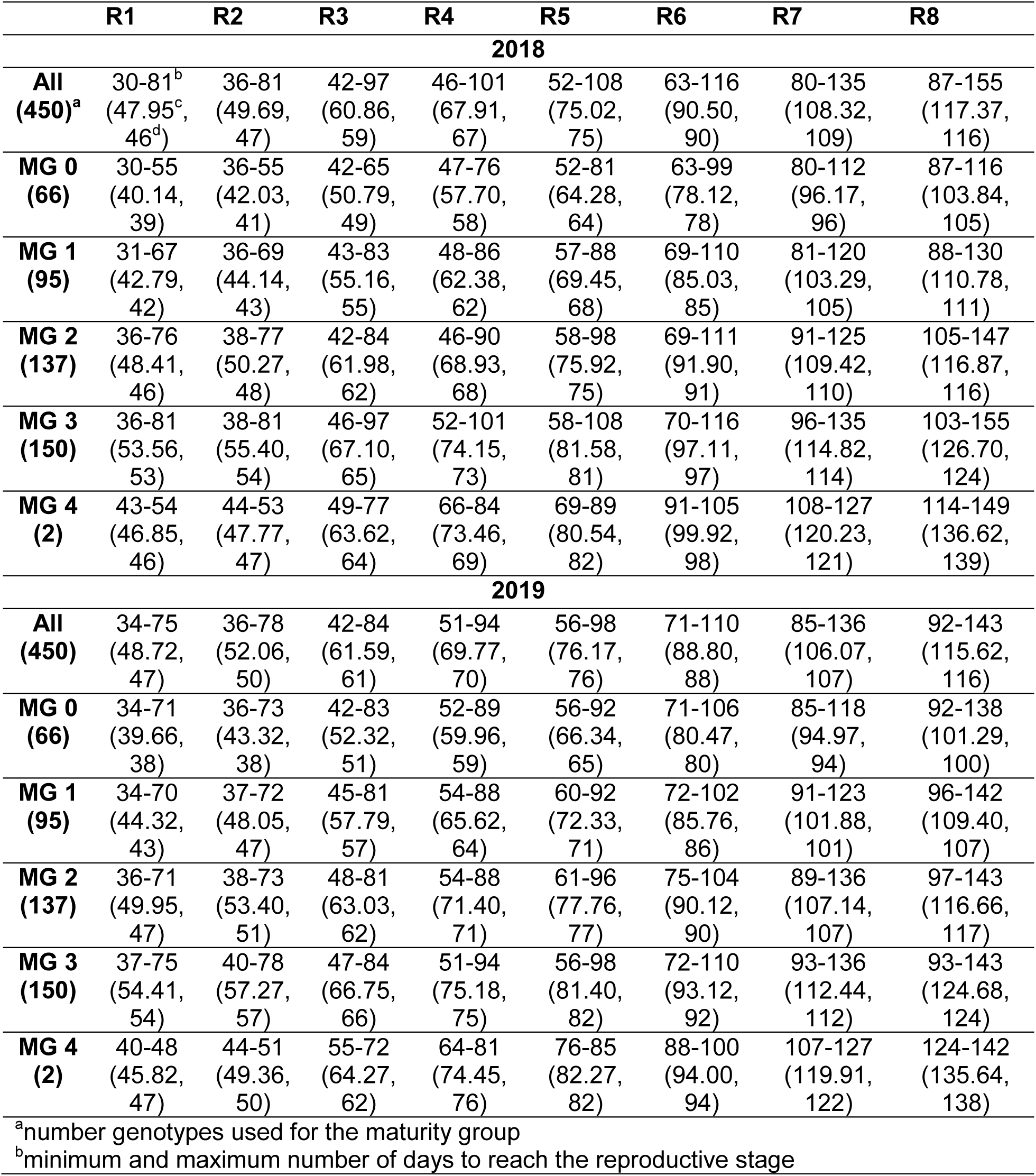
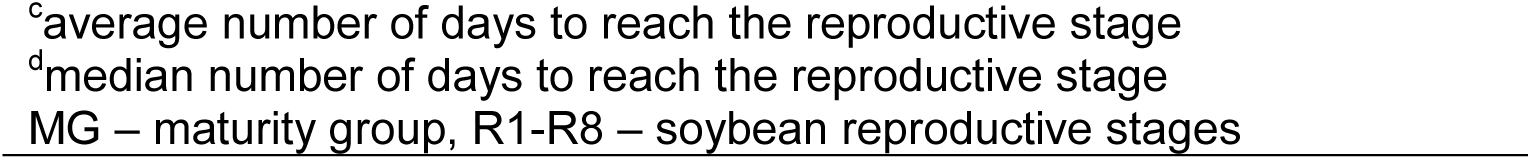
Days from planting to each reproductive development stage in both years.

|  | R1 | R2 | R3 | R4 | R5 | R6 | R7 | R8 |
| --- | --- | --- | --- | --- | --- | --- | --- | --- |
| **2018** |  |  |  |  |  |  |  |  |
| **All (450)<sup>a</sup>** | 30-81<sup>b</sup><br>(47.95<sup>c</sup>,<br>46<sup>d</sup>) | 36-81<br>(49.69,<br>47) | 42-97<br>(60.86,<br>59) | 46-101<br>(67.91,<br>67) | 52-108<br>(75.02,<br>75) | 63-116<br>(90.50,<br>90) | 80-135<br>(108.32,<br>109) | 87-155<br>(117.37,<br>116) |
| **MG 0 (66)** | 30-55<br>(40.14,<br>39) | 36-55<br>(42.03,<br>41) | 42-65<br>(50.79,<br>49) | 47-76<br>(57.70,<br>58) | 52-81<br>(64.28,<br>64) | 63-99<br>(78.12,<br>78) | 80-112<br>(96.17,<br>96) | 87-116<br>(103.84,<br>105) |
| **MG 1 (95)** | 31-67<br>(42.79,<br>42) | 36-69<br>(44.14,<br>43) | 43-83<br>(55.16,<br>55) | 48-86<br>(62.38,<br>62) | 57-88<br>(69.45,<br>68) | 69-110<br>(85.03,<br>85) | 81-120<br>(103.29,<br>105) | 88-130<br>(110.78,<br>111) |
| **MG 2 (137)** | 36-76<br>(48.41,<br>46) | 38-77<br>(50.27,<br>48) | 42-84<br>(61.98,<br>62) | 46-90<br>(68.93,<br>68) | 58-98<br>(75.92,<br>75) | 69-111<br>(91.90,<br>91) | 91-125<br>(109.42,<br>110) | 105-147<br>(116.87,<br>116) |
| **MG 3 (150)** | 36-81<br>(53.56,<br>53) | 38-81<br>(55.40,<br>54) | 46-97<br>(67.10,<br>65) | 52-101<br>(74.15,<br>73) | 58-108<br>(81.58,<br>81) | 70-116<br>(97.11,<br>97) | 96-135<br>(114.82,<br>114) | 103-155<br>(126.70,<br>124) |
| **MG 4 (2)** | 43-54<br>(46.85,<br>46) | 44-53<br>(47.77,<br>47) | 49-77<br>(63.62,<br>64) | 66-84<br>(73.46,<br>69) | 69-89<br>(80.54,<br>82) | 91-105<br>(99.92,<br>98) | 108-127<br>(120.23,<br>121) | 114-149<br>(136.62,<br>139) |
| **2019** |  |  |  |  |  |  |  |  |
| **All (450)** | 34-75<br>(48.72,<br>47) | 36-78<br>(52.06,<br>50) | 42-84<br>(61.59,<br>61) | 51-94<br>(69.77,<br>70) | 56-98<br>(76.17,<br>76) | 71-110<br>(88.80,<br>88) | 85-136<br>(106.07,<br>107) | 92-143<br>(115.62,<br>116) |
| **MG 0 (66)** | 34-71<br>(39.66,<br>38) | 36-73<br>(43.32,<br>38) | 42-83<br>(52.32,<br>51) | 52-89<br>(59.96,<br>59) | 56-92<br>(66.34,<br>65) | 71-106<br>(80.47,<br>80) | 85-118<br>(94.97,<br>94) | 92-138<br>(101.29,<br>100) |
| **MG 1 (95)** | 34-70<br>(44.32,<br>43) | 37-72<br>(48.05,<br>47) | 45-81<br>(57.79,<br>57) | 54-88<br>(65.62,<br>64) | 60-92<br>(72.33,<br>71) | 72-102<br>(85.76,<br>86) | 91-123<br>(101.88,<br>101) | 96-142<br>(109.40,<br>107) |
| **MG 2 (137)** | 36-71<br>(49.95,<br>47) | 38-73<br>(53.40,<br>51) | 48-81<br>(63.03,<br>62) | 54-88<br>(71.40,<br>71) | 61-96<br>(77.76,<br>77) | 75-104<br>(90.12,<br>90) | 89-136<br>(107.14,<br>107) | 97-143<br>(116.66,<br>117) |
| **MG 3 (150)** | 37-75<br>(54.41,<br>54) | 40-78<br>(57.27,<br>57) | 47-84<br>(66.75,<br>66) | 51-94<br>(75.18,<br>75) | 56-98<br>(81.40,<br>82) | 72-110<br>(93.12,<br>92) | 93-136<br>(112.44,<br>112) | 93-143<br>(124.68,<br>124) |
| **MG 4 (2)** | 40-48<br>(45.82,<br>47) | 44-51<br>(49.36,<br>50) | 55-72<br>(64.27,<br>62) | 64-81<br>(74.45,<br>76) | 76-85<br>(82.27,<br>82) | 88-100<br>(94.00,<br>94) | 107-127<br>(119.91,<br>122) | 124-142<br>(135.64,<br>138) |

<sup>a</sup>number genotypes used for the maturity group
<sup>b</sup>minimum and maximum number of days to reach the reproductive stage
<sup>c</sup>average number of days to reach the reproductive stage <sup>d</sup>median number of days to reach the reproductive stage MG – maturity group, R1-R8 – soybean reproductive stages

### 3.2 Effect of environmental variables and genotype-environment (G x E) interaction

We evaluated correlations between reproductive phases and four environmental variables: DL, DTR, GDD, and PTT to investigate how environmental cues influence reproductive development (Figure 2A, Supplementary File 1 - Figure S2). The reproductive stages showed a strong correlation with DL and GDD (Figure 2B, Supplementary File 2 - Table S7). Environmental factors also had a significant impact on transition times, albeit the effects varied depending on the kind of environment (Supplementary File 1 - Figure S3). Based on these results, DL and GDD were identified as important regulators as they represent two separate biological mechanisms, such as photoperiod and thermal time, that control flowering and reproductive development.

**Figure 2.**
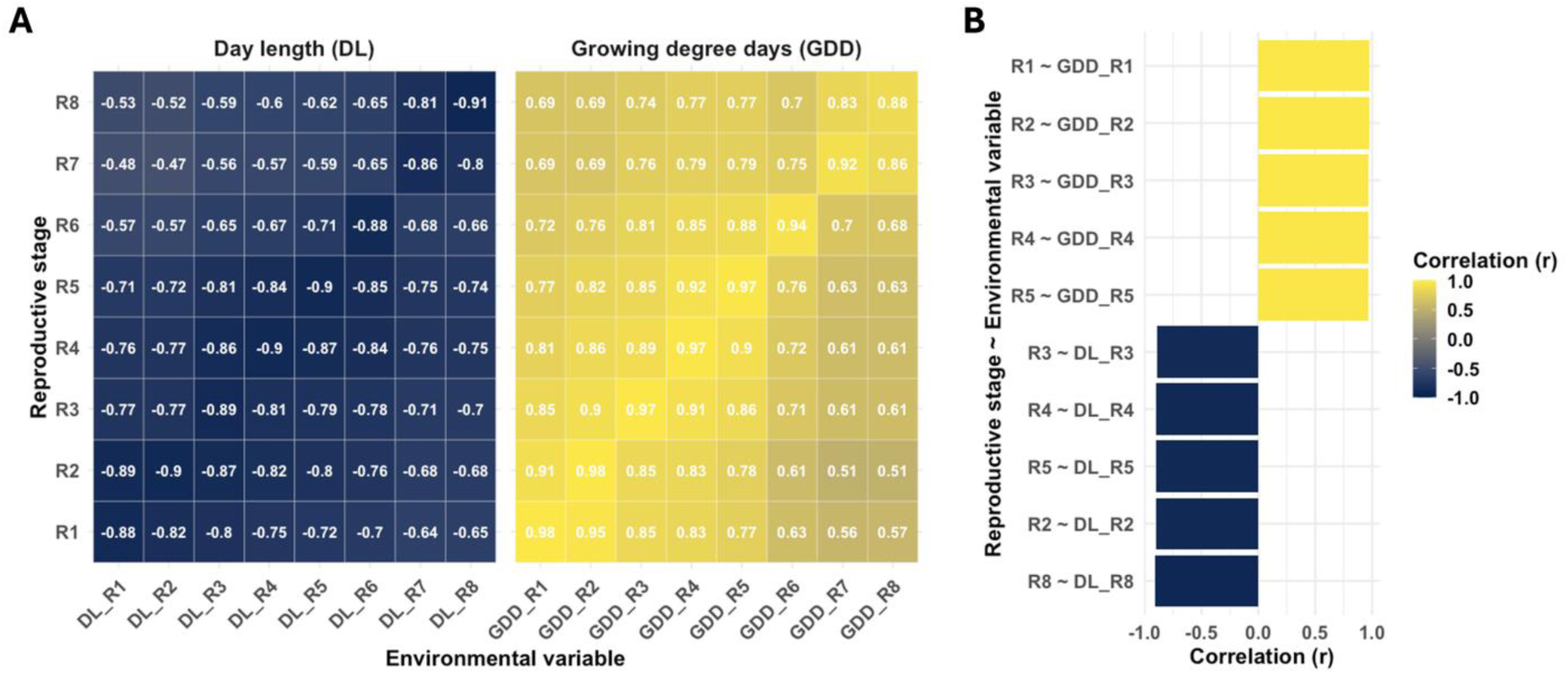
Correlation between reproductive stages and environmental variables in a soybean diversity panel. (A) Pearson correlation coefficients for environmental variables such as day length (DL), and growth degree days (GDD) with reproductive stages (R1– R8), (B) Top ten correlations between environmental variables and reproductive stages.

Genotype and G x E interaction effects significantly contributed to the observed phenotypic variability for reproductive stages and their transition traits. The DL and GDD showed a wide range of environment-adjusted means for each reproductive stage (R1– R8). The VCA revealed that genotype effects accounted for the lowest phenotypic variation at R6 (66%) to the highest at R1 (81%), with the G x E interaction accounting for an additional 6.81% at R1 to 8.08% at R6, while replication and residual variance proportions were quite minor (Supplementary File 2 - Table S8). The developmental trajectory of most genotypes for both DL and GDD models was parallel or slightly divergent, suggesting modest plasticity and G x E interaction (Supplementary File 1 - Figure S4A and S4B). The average responses at R1 and R2 for DL (0.040 and 0.055, respectively) were marginally stronger than those for GDD (0.049 and 0.056, respectively). However, the GDD model revealed significantly higher plasticity in the mid-reproductive stages (R3-R5): mean slopes at R3 and R5 were 0.029 and 0.026, respectively. For DL, mean slopes were higher: mean slopes at R3 and R5 were 0.034 and 0.028, respectively. At R8, DL had similar slope as GDD (DL mean slope = 0.026; GDD = 0.027) (Supplementary File 2 - Table S9). Genotype effects for reproduction transitions accounted for an average of about 33.3% of the total variance, with an additional 12.4% explained by the G x E interaction (Supplementary File 2 - Table S10). Reaction norm analysis of transition traits exhibited significant variance in the flexibility of genotypes throughout developmental intervals. According to mean absolute slope values, the first transition (R1R2) had the highest flexibility, and DL-based models captured somewhat larger average slopes (0.3225) than GDD-based models (0.3054). The mid-stage transitions (such as R2R6, R4R6, and R5R6) responded better to GDD, indicating that thermal accumulation is a key factor in controlling these transitions. Slope values remained high for later transitions (R6R7–R7R8), but variation increased, especially for DL, suggesting greater sensitivity to coupled environmental signals or genotypic variability (Supplementary File 2 - Table S11).

### 3.3 Effect of genomic loci during reproductive stages and their transitions

We conducted the GWAS using BLUE estimates from both DL and GDD models for the reproductive stages and their transitions. Two SNPs encoding the *Dt1* gene (Glyma.19G194300) and *E2* gene (Glyma.10G221500) were consistently detected at chromosome 19 and chromosome 10, respectively, for both DL and GDD (Supplementary File 1 - Figure S5A & S5B, and Supplementary File 2 - Table S12 & S13). The *Dt1* gene was detected for the mid reproductive stages, such as R4-R6, while the *E2* gene was found to be associated with later maturity stages, R7 and R8.

GWAS using two reaction norm parameters, plasticity slope and intercept, confirmed the importance of *Dt1*. Using slope, the *Dt1* gene was detected for the R5 stage using the DL data across genotypes. *E2* gene was not detected for any stage using the DL data. However, we identified SNPs for the *E1* gene (*Glyma.06G207800*) on chromosome 6 for reproductive stage R2, R6, and R8 (Supplementary File 1 - Figure S5C, Supplementary File 2 - Table S12 & S13). The *E1* gene was also detected for the R2 stage as well as when intercept was used as a response variable using the GDD data. None of the known maturity-related genes was detected for both plasticity slope and intercept. However, we found one QTL from DL data at position Chr8: 17603027 (*Glyma.08G216800*) associated with plasticity slope and intercept for R3 stage, as well as BLUE and plasticity slope for R1 stage. This QTL was identified as PPPDE putative thiol peptidase family protein (Supplementary File 2 - Table S12). For GDD, we also identified one SNP at position Chr4:7556514 associated with plasticity slope and intercept for the R7 stage. Using Soybase genome browse, we found that the nearest (within 1Kbp) gene, annotated as *Glyma.04G088100*, which is an RNA-binding KH domain-containing protein. In case of reproductive stage transitions using DL, we identified the *Dt1* and *Dt2* genes on chromosomes 19 and 18, respectively, and the *E1* gene on chromosome 6 (Supplementary File 2 - Table S14). For GDD, we also detected the *Dt1* and *Dt2* genes on chromosomes 19 and 18, respectively, and the *E1* and *E2* genes were detected on chromosomes 6 and 10, respectively (Supplementary File 2 - Table S15).

Besides the above-mentioned maturity-related genes, several high-confidence SNPs were detected from multiple GWAS methods. Among them, most of the SNPs were unknown or uncharacterized. Considering the DL environmental gradient, a few top SNPs that were detected through three or four GWAS models were: Chr01:51691145 (*Glyma.01G180600*), Tetratricopeptide repeat (TPR)-like superfamily protein (BLUE: R1, R2, R3, R4, R5, R6; Intercept: R2, R3, R4); Chr06:3396872 (*Glyma.06G044600*), NAD(P)-binding Rossmann-fold superfamily protein (Slope: R1, R3, R5, R7); Chr08:1017668 (*Glyma.08G013000*), Myosin heavy chain-related protein (Slope: R4); and Chr10:45269968 (*Glyma.10G221300*), S-adenosylmethionine carrier 1 (BLUE: R7, R8; Intercept: R1, R5, R6, R7, R8). The SNP at position Chr01:51691145 was also detected for GDD through three or four GWAS models, and the SNP at Chr10:45269968 was found to be strongly connected with the *E2* gene at the haplotype network.

Haplotype blocks were built around each GWAS-significant SNP based on SNPs in high linkage disequilibrium (LD ≥ 0.8) to understand the genetic structure and linkage patterns of detected maturity genes. The analysis of six GWAS SNPs revealed 68 distinct haplotypes, indicating a significant level of allelic variation within genomic segments characterized by LD (Supplementary File 2 - Table S16). LD correlation (r) values ranged from 0.91 to 1.0, and a mean r value of 0.96. Ten LD blocks were found for the top five GWAS hits, highlighting their potential as key hubs in the design of haplotype networks (Supplementary File 1 - Figure S6, and Supplementary File 2 - Table S17). Compared to single SNPs, these haplotype blocks offer a finer-scale resolution for association mapping and might be better at capturing causal variation. Strong LD and significant haplotype diversity point to strong genomic areas that need more functional investigation, especially considering the observed trait variability.

### 3.4 Random Forest-based maturity model dissecting genetic and environmental contributions to soybean phenology

Random forest (RF) models trained on the NIL and diversity panels exhibited high predictive power across soybean reproductive stages (R1 to R8), with some variability observed between early and late stages. With RMSE values ranging from 0.01 to 1.98 and corresponding R^2^ values between 0.96 and 0.99, the model with NIL panel performed well for all reproductive phases except R8 (Supplementary File 1 - Table S3). On the other hand, the same model using the diversity panel demonstrated consistently better accuracy at every level, including R8. Different patterns of environmental variable relevance over phases were found by the feature importance analysis based on the percentage increase in mean squared error (MSE) (Figure 3A and Figure 4A). Environmental predictors were the main factors affecting stage transitions in both panels, especially GDD and DL. In particular, GDD at R1 and R2 stages continuously received the greatest relevance ratings during early stages (R1– R3) in both panels. As we moved toward a more complex environmental integration, both GDD and DL from prior stages (e.g., GDD_R4, DL_R5) helped to explain variance for intermediate stages (R4–R6). The diversity panel indicated that DL_R8 and GDD_R8 were predominantly dominant for late stages (R7–R8), whereas the NIL panel emphasized DL_R2 and DL_R7 (Figure 3A and Figure 4A). Throughout most phases, maturity-related genes (*E1–E4*, *Dt1*, *Dt2*) made a minor contribution to the R8 stage prediction, particularly in the diversity panel.

**Figure 3.**
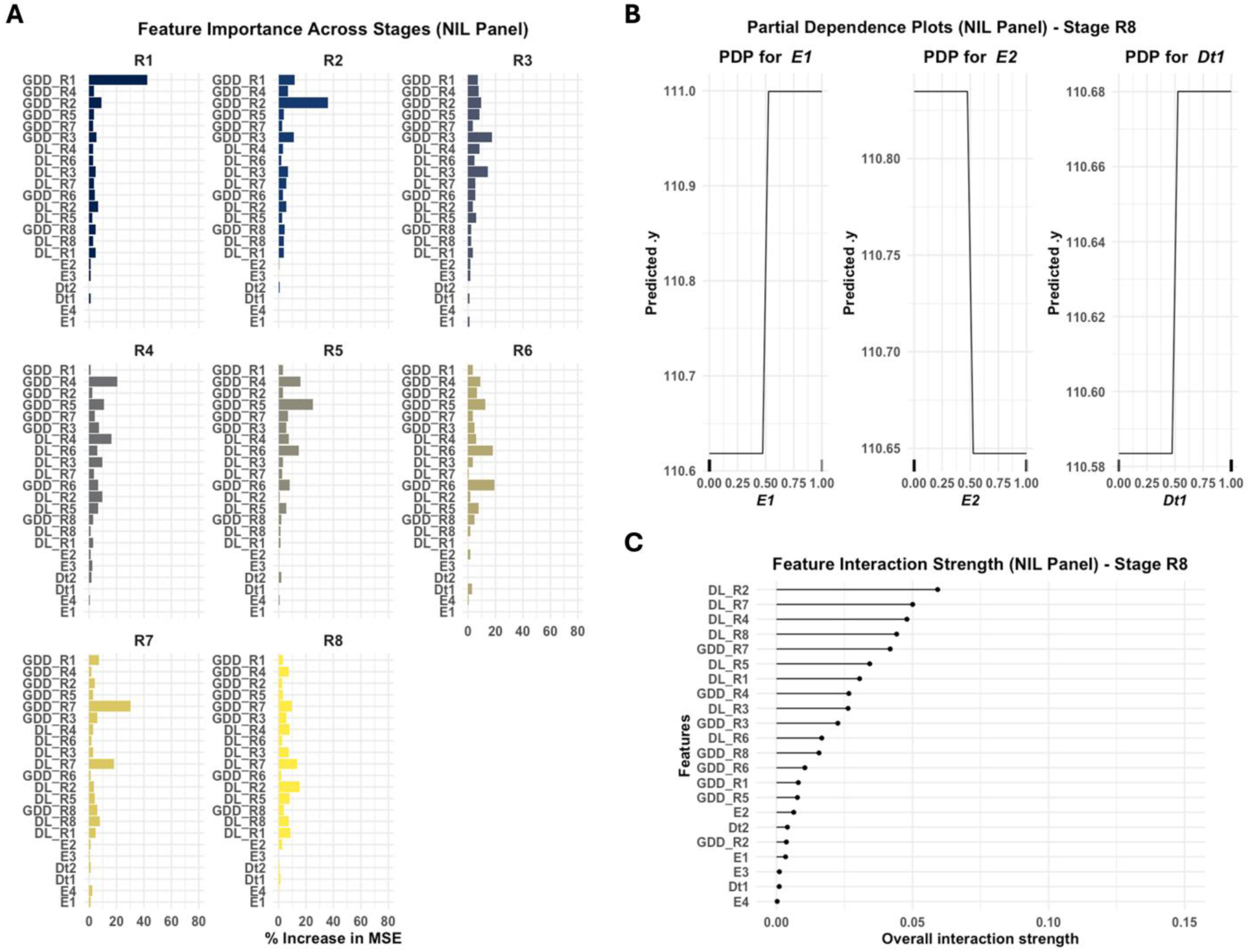
Random Forest-based maturity model of the reproductive stages from NIL population. (A) Feature importance by stage as a percentage increase in mean square error (MSE) for reproductive stages R1 through R8, (B) One-dimensional partial dependence plot (PDP) that illustrates how *E1*, *E2*, and *Dt1* marginally affect the expected R8 values, and (C) Analysis of feature interaction strength for stage R8. DL and GDD indicate day length and growing degree days, respectively.

**Figure 4.**
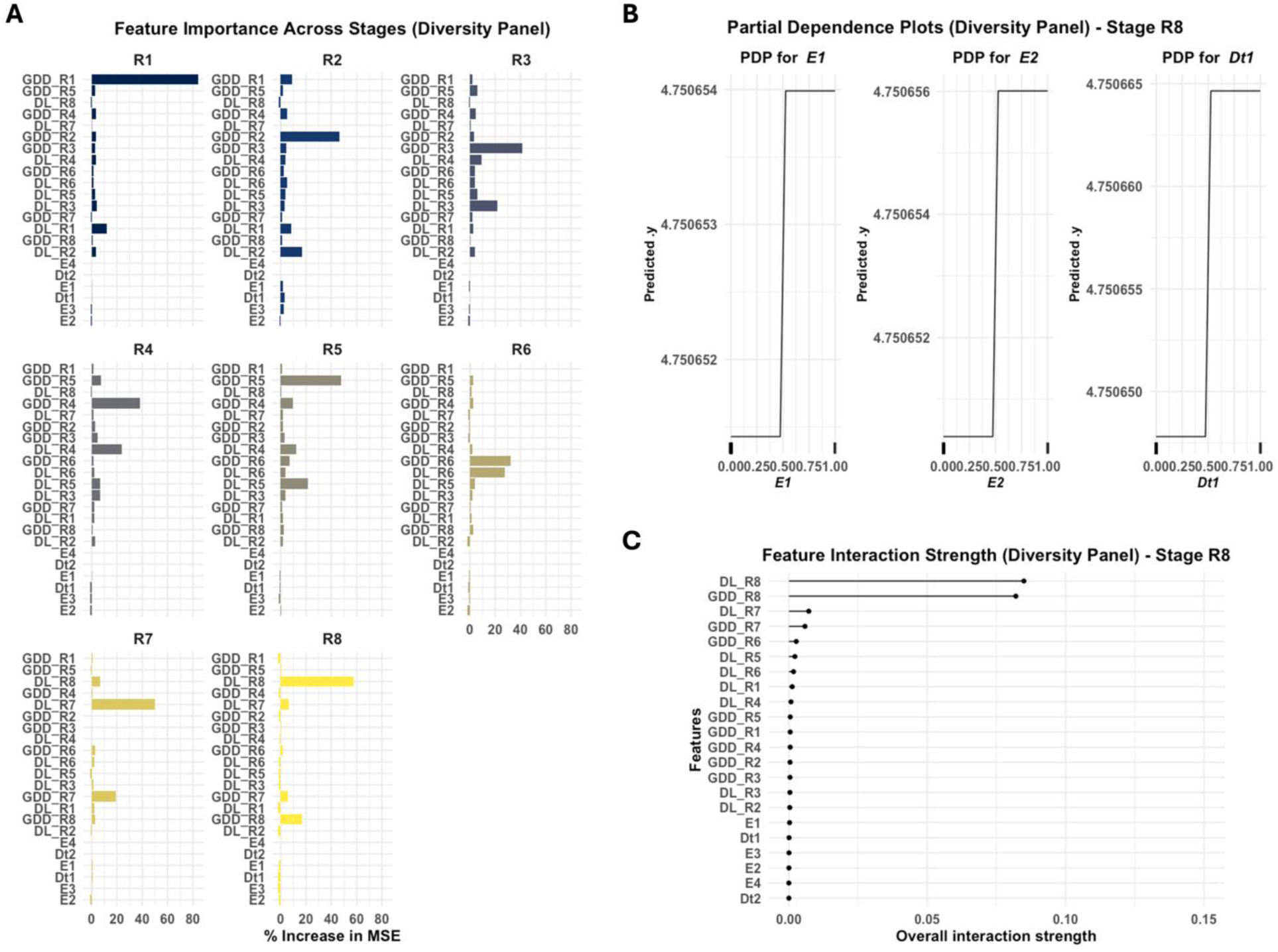
Random Forest-based maturity model of the reproductive stages from the diversity panel. (A) Feature importance of the variables used in the model for reproductive stages as a percentage increase in mean square error (MSE), (B) One-dimensional partial dependence plot (PDP) showing the effect of *E1*, *E2*, and *Dt1* gene on R8 stage, and (C) Feature interaction strength for R8 stage. DL and GDD indicate day length and growing degree days, respectively.

The RF model also provided information regarding specific developmental stages. For instance, at growth stage R8, the partial dependency plots showed that *E1*, *E2*, and *Dt1* genes had distinct stepwise effects in both panels (Figure 4B and Figure 4B). However, overall marginal effects on the anticipated R8 value were small, which is consistent with their low feature selection relevance rankings. Figures 3C and 4C show the overall interaction contributions of each variable used in the model as measured by the interaction strength analysis. The NIL panel’s most interactive features were DL at the R2 and R7 stages, followed by DL at the R3 and R5 stages, indicative of NIL’s sensitivity to cross-stage photoperiods. Particularly in the diversity panel, where DL and GDD at the R8 stage were predominant drivers of interaction effects, genetic factors (*E*-loci) exhibited the least amount of interaction strength, but environmental predictors (DL and GDD) contributed the most to interaction terms.

## 4. DISCUSSION

In this study, we presented results from a developmental perspective of dissecting phenotypic variation in reaching each of the reproductive stages under natural field conditions, considering environmental gradients, and providing insight into the genetic loci regulated by the effect of genotype, environmental variable, and their interaction. We provide the first comprehensive stage interval study from R1 to R8, elucidating novel insights. Finally, we developed a random forest-based model that can predict the effects of genotype and environmental variables in the reproductive stages of soybean.

Using two years of data and a diversified soybean panel consisting of >450 PI accessions and >65 NILs, our study showed significant phenotypic variation in reproductive stages and their transitions. The change in the environment highlights how day length has a significant impact on the early stages of reproductive development in soybean. These kinds of variations due to genetic and non-genetic factors were also reported previously (52,53). These findings highlight that, even for the reproductive stages under strong genetic control, their expression is context-dependent. Environmental factors become the main determinants of reproductive development. The temperature and photoperiod control flowering and maturity through mostly separate but temporally synchronized processes (14,54,55). In this study, GDD accounted for more variance in mid-to-late reproductive stages, which is consistent with the requirement for thermal accumulation for pod filling and maturity, whereas DL had a significant influence on early transitions (Figure 2A). Early transitions are driven by photoperiod, where short DL triggers early flowering while long DL delayed flowering (56,57). G x E occurs frequently, is linked to a variety of genetic factors and molecular events, and is frequently brought about by changes in the strength of genetic effects in response to environmental stimuli (58). In our study, we found stage-specific contributions of genotype, environmental variables, and G x E interaction effects. For instance, early stages like R1 displayed the highest genetic effect, which suggests that early floral initiation is more genetically controlled than the start of pod filling. Early stages were more receptive to DL, while mid stages are governed by GDD or heat accumulation that is related to pod setting and seed development, and later maturity is tandemly influenced by both cues. These kinds of dynamic environmental influences on reproductive trajectories were also reported by earlier studies (59,60), which have shown that this sensitivity to temperature and photoperiod varies between different maturity groups and genetic origins. Photoperiod sensitivity is often higher in later maturity groups, and high temperatures can either increase or decrease these sensitivities (61). The cumulative impacts of earlier and later developmental stimuli were also incorporated in transition stages (e.g., R1R8, R2R8). When compared to individual reproductive stages, the transition stages showed larger G x E and residual variance, indicating that the impact of the environment increased when considering reproductive transition stages. With the large genetic panel and reproductive stages and their transitions, these findings further strengthen our understanding of the genotypic effect and its interaction with the environment, both for reproductive stages and their transitions. However, genetics is one of the main factors influencing growth stages and their intervals.

GWAS from this study identified key loci, including *Dt1*, *E1*, and *E2,* for different reproductive stages and their transitions (Supplementary File 1 - Figure S5), which were also detected in a meta-GWAS study and known to regulate flowering and maturity (62). Recent research showed that *E1Lb*, independent of *E1*, delays flowering in long-day circumstances (63). The *E2* gene, orthologous to the *GIGANTEA* (GI) gene in *Arabidopsis thaliana*, was found to be involved in delaying flowering by inhibiting the expression of GmFT2a during long days (26). The *Dt1* gene is involved in the determination of plant growth type, such as determinate or indeterminate type (64). These genes’ pleiotropic roles across reproductive phases were confirmed by the detection of these genes not only for reproductive stages but also for stage transitions. The findings of the study conducted by Miranda et al. (2020) enabled the comparison of various variant alleles of those genes and showed notable genotype-based differences between days to flowering and days to maturity (34). We report novel loci, including *Glyma.08G216800* (PPPDE putative thiol peptidase family protein) and *Glyma.04G088100* (RNA-binding KH domain-containing protein) from GWAS with slope and intercept of the reaction norm, which revealed more levels of regulatory intricacy that may be connected to environmental responsiveness, which varies by reproductive stage. The PPPDE putative thiol peptidase family protein was identified for early reproductive stages, indicating that, irrespective of genotype and environmental effect, it might have a regulatory role with photoperiod during the early reproductive stage. But its functions are yet unclear and need additional research. It has been reported that the PPPDE superfamily is a deubiquitinase (DUB) that is conserved in eukaryotes, including humans (65). Arabidopsis *AtC3H59* controls cell division by interacting with the PPPDE family protein Desi1 via its WD40 domain, and the PPPDE domain of the deubiquitinase PICI1 serves as an immune hub for pattern-triggered immunity and effector-triggered immunity in rice (66). We also identified the RNA-binding K homology (KH) domain-containing protein associated with the late reproductive stage, suggesting that this protein might be regulated by thermal or heat accumulation of plants rather than by the genotype effect. Guan et al. (2013) reported that RCF3 is a nuclear-localized putative RNA-binding protein that contains a KH domain. The *rcf3 Arabidopsis* mutant was more resistant to heat stress than the wild-type, which is consistent with the general higher accumulation of heat-responsive genes (67). However, in our study, we did not investigate the effect of temperature on the reproductive stages, and we did not functionally validate the RNA-binding K homology (KH) domain-containing protein. The novel SNP at position Chr01: 51691145 (*Glyma.01G180600*), which is a tetratricopeptide repeat (TPR)-like superfamily protein detected from BLUE and intercept for both DL and GDD for early to mid-reproductive stages, indicated that this protein could be a potential target for genetic regulation of reproductive stages. This TPR protein has been reported to be involved in hormonal regulation and drought stress in plants (68,69). In the haplotype network, the *E2* gene or the GIGANTEA protein was closely connected with NOD26-like intrinsic protein, Nucleotide-sugar transporter family protein, Ribosomal protein S13/S18 family, S-adenosylmethionine carrier 1, Inositol polyphosphate 5-phosphate-related protein, and five other uncharacterized or unknown proteins. Among the uncharacterized or unknown proteins, *Glyma.10G221900* had the highest LD with the *E2* gene (Supplementary File 1 - Figure S6). According to the reported function of the above genes that were in high LD with the *E2* gene, they are mainly involved in the growth and development of plants in several cellular pathways (70–78). However, for better understanding of the mechanism of development during maturity could be possible by discovering the function of unknown or uncharacterized proteins found in strong LD and varied haplotype structures surrounding GWAS-significant SNPs.

The Random Forest models corroborated experimental findings by attributing dominant predictive power to environmental variables, especially DL and GDD, with minor contributions from known maturity genes (Figure 3 and 4). The difference in prediction accuracy between the diversity and NIL panels demonstrates how more genetic variation improves the generalizability of the model. The improved model performance of the diversity panel could be attributed to increased resolution and variability in genotype and environmental responses, which would enable the model to more effectively partition and learn intricate patterns influencing phenology. Analysis of feature importance and interaction strength showed a temporal pattern of cue dominance. It’s interesting to note that even well-known maturity genes had negligible effect sizes in prediction models, confirming the idea that the environment mostly controls the timing of reproductive development in a diverse population. Since this study didn’t include commercial varieties, it is difficult to generalize these findings to advanced breeding stock. However, the large genetic collection and NILs were very useful to develop a prediction model that can be utilized to predict stage-specific regulation of genotypic and environmental factors with applications in breeding programs. These discoveries open the door to more focused breeding tactics that take advantage of developmental plasticity and environmental sensitivity. For example, breeders can leverage this information for ideotype breeding to maximize crop productivity by optimizing reproductive stages and their transitions in conjunction with physiological characteristics in cultivation regions (6). The crop modelers can utilize the data and results presented in this study for more rigorous crop modeling applications. However, a common approach for such studies is to perform multi-year studies at the same geographical location to minimize the effect of photoperiod, and to some extent, temperature or thermal accumulation. While less comprehensive and a more limited scope of results interpretation and extrapolation than an extensive study across multiple years, locations, and geographies, it has advantages due to better and more standardized experimental factors. Also, central Iowa in the U.S. falls in primarily the MGII maturity group, which has among the highest soybean production in the U.S. and Canada (13,79).

## 5. CONCLUSION

From a comprehensive viewpoint based on the results of phenotypic and genomic analysis and statistical modeling using a large panel of genotypes, the study demonstrated that intricate interactions between genotype, environment, and their interaction drive the reproductive development of soybeans. This study provides practical insights for breeders looking to optimize soybean maturity and yield under varying environmental gradients for applying ideotype breeding concepts. Additionally, farmers can make better decisions about variety selection by identifying stage-specific environmental sensitivity, where photoperiod (DL) mostly influences early and late reproductive stages and thermal time (GDD) primarily influences mid-stages. Growers can lessen the chance of yield loss during crucial reproductive times due to heat, drought, or frost stress by coordinating reproductive transitions with favorable photothermal windows. The identification of novel loci governing plasticity and the major maturity genes (*E1*, *E2*, *Dt1*, and novel candidate genes identified in this study) gives plant breeders a genetic toolbox for creating varieties with specific adaption. Breeders can find genotypes with advantageous environmental responsiveness under a variety of situations by combining BLUE-based genetic value estimates, reaction norm modeling, GWAS, and a random forest-based predictive model. When taken as a whole, these results lend credence to breeding plans that try to increase phenological plasticity and give farmers the ability to match genotypes to certain environmental profiles, which eventually improves output stability and resource efficiency. However, our study was limited to central Iowa only and was conducted for two years. Multi-year data from multiple geographical locations could provide more rigorous results of the effect of environmental gradients and G x E interaction. To speed up breeding for environment-specific maturity optimization, future studies should concentrate on the functional validation of novel loci and the mechanistic analysis of genotype-by-environment responsiveness.

## Data availability

Upon the publication of this article, all data related to this study will be available at https://github.com/SoylabSingh/SoyMaturity

## Author Contributions

Conceptualization: SC, JMS, AKS; Data curation: JMS; Methodology: SC, JMS, AKS; Formal data analysis and visualization: SC, JMS; Writing original draft: SC; Review and editing: SC, AKS; Project administration, funding acquisition, resources, and supervision: AKS.

## Acknowledgements

1. J. Shook was partially supported by funding from the NSF-NRT. Funding support came from R. F. Baker Center for Plant Breeding, Iowa Soybean Association, USDA-NIFA, and National Science Foundation. Research support from members of AKS group at ISU - staff, post-doctoral fellows, and students - is sincerely acknowledged.

